# Improved Version of ChETA Promotes Aggression in the Medial Amygdala

**DOI:** 10.1101/2022.06.05.493862

**Authors:** Rongfeng K. Hu, Patrick B. Chen, André Berndt, David J. Anderson, Weizhe Hong

**Affiliations:** Department of Neurobiology and Department of Biological Chemistry, University of California, Los Angeles, CA 90095, USA; Institute for Stem Cell and Regenerative Medicine, Department of Bioengineering, University of Washington, Seattle, WA 98109, USA; Division of Biology and Biological Engineering, Tianqiao and Chrissy Chen Institute for Neuroscience, and Howard Hughes Medical Institute, California Institute of Technology, Pasadena, CA 91125, USA; Institute for Translational Brain Research and Zhongshan Hospital, Fudan University, Shanghai, 200032, China

## Abstract

The development of optogenetic tools has significantly advanced our understanding of neural circuits and behavior. The medial amygdala, posterior dorsal subdivision (MeApd) is part of a distributed network controlling social behaviors such as mating and aggression. Previous work showed that activation of GABAergic neurons in mouse MeApd using channelrodopsin-2 (ChR2^H134R^) promoted aggression. In a recent study, Baleisyte et al. (2022) confirmed these findings using the same reagents (i.e. ChR2^H134R^), but also reported that a different ChR2 variant with faster kinetics—ChETA—inhibited rather than promoted aggression when high laser power, long duration photostimulation conditions were used. As ChETA is known to have a substantially lower photocurrent than ChR2 and other opsins, an improved version of ChETA (i.e. ChR2^E123T/T159C^; ChETA_TC_) was subsequently developed. ChETA_TC_ has larger photocurrents than the original ChETA while maintaining fast kinetics and low plateau depolarization. Here we show that activating MeApd GABAergic neurons using the improved ChETA_TC_ promotes aggression, similar to ChR2^H134R^, suggesting that the results obtained using the original ChETA are not due to a difference in channel kinetics. Furthermore, we found that ChETA_TC_ is capable of driving a rapid onset of aggression within 200-300 milliseconds of stimulation, suggesting that this effect reflects direct activation of MeApd GABAergic neurons. We conclude that the different behavioral phenotypes observed using the original ChETA vs. ChETA_TC_ and ChR2 likely reflects the weaker photocurrents in ChETA vs. other opsins, and/or the long duration/high power photostimulation conditions used with ChETA. Consistent with this conclusion, the results obtained using ChR2 or ChETA_TC_ are complementary to findings from loss-of-functions experiments using optogenetic inhibition, chemogenetic inhibition, and neuronal ablation. These data support a positive-acting role of MeApd Vgat^+^ neurons in aggression. Our findings, in conjunction with studies of Berndt et al. (2011), suggest that the improved ChETA_TC_ should be used when faster kinetics than ChR2 offers are required.

## Main text

Optogenetics has transformed research on neural circuits and behavior by enabling precise manipulation of neural activity with high spatiotemporal resolution (Deisseroth, 2015; Fenno et al., 2011; Kim et al., 2017). Over the last two decades, a large array of opsins with different properties have been developed for a variety of applications (Mattis et al., 2012; Zhang et al., 2011). Channelrodopsin-2 (ChR2^H134R^; hereafter referred to simply as ChR2) was one of the first opsins developed to activate neurons (and nonneuronal cells) and has been widely used in neuroscience research (Fig. 1A). Since then, many new variants of ChR2 have been developed for various applications. Among them, ChETA (ChR2^E123T/H134R^) is a variant that displays fast kinetics to facilitate the study of fast-spiking neurons which require stimulation at 40 Hz or above (Gunaydin et al., 2010). Although ChETA exhibits fast kinetics, it appears to have a substantially lower photocurrent than ChR2 (Baleisyte et al., 2022; Berndt et al., 2011; Mattis et al., 2012). Consequently, very few published studies have used the original ChETA, in comparison to those using ChR2 (Figure 1A). To address this problem, an improved version of ChETA (ChR2^E123T/T159C^; ChETA_TC_) was subsequently developed (Berndt et al., 2011; Mattis et al., 2012). This improved version carries an additional mutation (T159C) besides the original ChETA mutation (E123T). ChETA_TC_ possesses ∼3-fold larger photocurrents than the original ChETA, while maintaining fast kinetics and low plateau depolarization (Fig. 1B; Berndt et al., 2011; Mattis et al., 2012).

**Figure 1.**
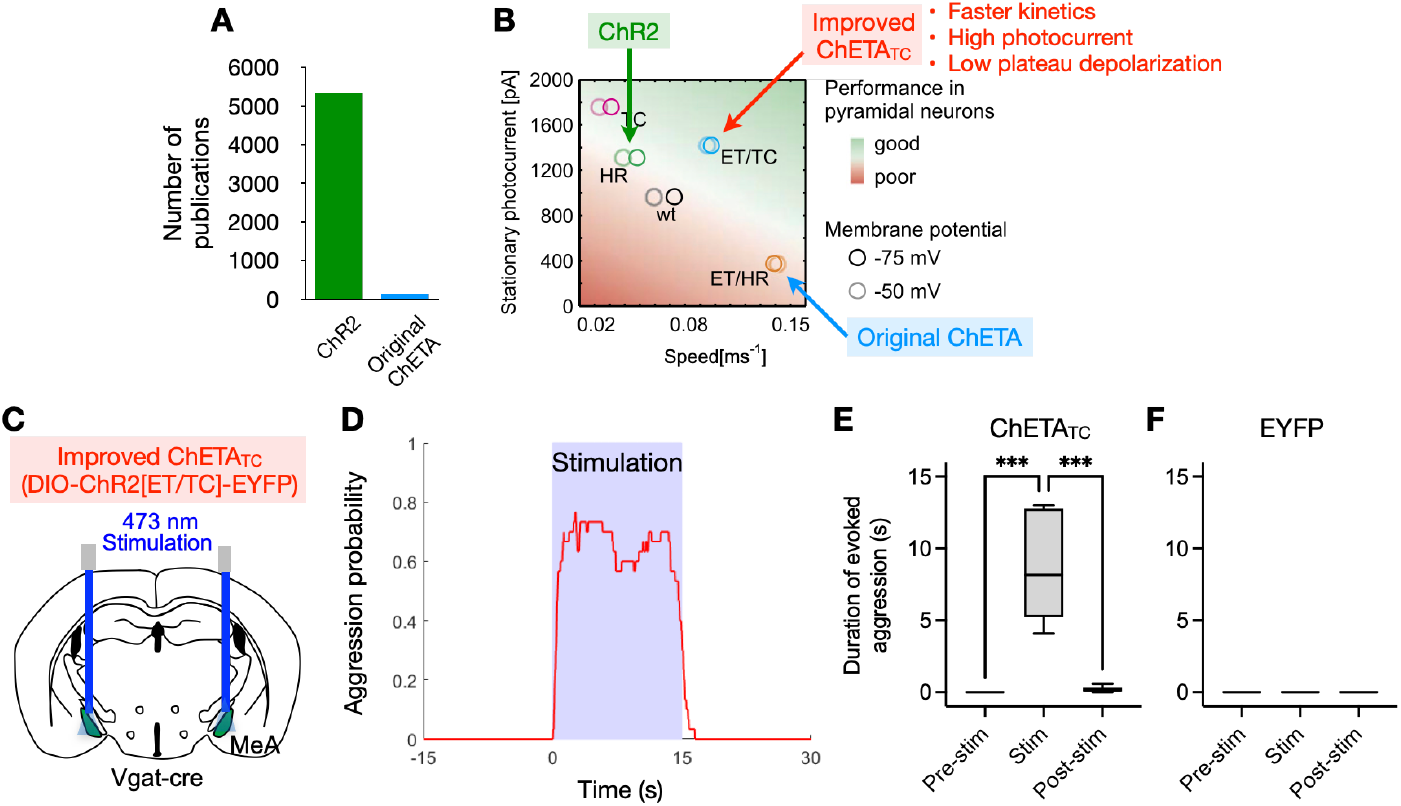
Improved ChETA_TC_ activation of MeA GABAergic neurons induces rapid, robust aggression. **A**, Number of publications that include ChR2 or the original ChETA on NCBI PubMed. **B**, Electrophysiological properties of various ChR2 variants. The improved ChETA_TC_ possesses higher photocurrents than ChETA while maintaining the fast kinetics and low plateau depolarization (adapted from Berndt et al 2011). **C**, Experimental design for ChETA_TC_ activation of MeA GABAergic neurons. **D**, Probability of aggression during photoactivation trials of MeA GABAergic neurons expressing ChETA_TC_. Each stimulation: 15 s of 20 Hz, 20 ms, 473 nm, ∼5 mW/mm^2^. **E, F**, Increased duration of aggression during photoactivation epochs (0–15s) relative to pre- (−15–0s) and post-stimulation (15– 30s) epochs. **E**, ChETA_TC_; **F**, EYFP control. One-way ANOVA with post-hoc Tukey’s multiple comparisons. *** *p* < 0.001. n = 5 animals for ChETA_TC_ and 4 animals for EYFP.

Using ChR2, we have previously shown that activation of GABAergic (Vgat^+^) neurons in the MeApd promoted aggression (Hong et al., 2014). Conversely, optogenetic silencing of MeApd GABAergic neurons (using eNpHR3.0) acutely suppressed ongoing aggression. These bi-directional gain- and loss-of-function phenotypes together suggested that MeApd GABAergic neurons exert a net positive-acting role in aggression. Consistent with these findings, ablation or chemogenetic silencing of aromatase^+^ GABAergic neurons in the MeApd decreased aggressive behavior (Unger et al., 2015), while chemogenetic activation of Npy1R+ neurons (predominantly GABAergic) in the MeA also promoted aggression, via projections to the BNST (Padilla et al., 2016; Unger et al., 2015). These studies further suggested that aggression can be promoted by specific subpopulations of GABAergic neurons within the MeApd.

That chemogenetic inhibition, neuronal ablation, and optogenetic inhibition of MeApd GABAergic neurons all suppressed aggression, while optogenetic activation of this population enhanced aggression, suggests a positive-acting role of MeApd GABAergic neurons in the control of aggression (Chen and Hong, 2018; Lischinsky and Lin, 2020; Raam and Hong, 2021). Importantly, the findings that the MeApd contains GABAergic neurons that promote aggression does not mean that all GABAergic neurons in MeApd promote aggression (Raam and Hong, 2021). Indeed, weaker activation of MeApd neurons with ChR2 using lower photostimulation intensities promoted mounting (likely to represent a weak form of agonistic behavior) rather than aggression (Hong et al., 2014; Karigo et al., 2021). This suggests that MeA GABAergic neurons can promote multiple types of social behaviors. Subsequent studies have shown that MeApd GABAergic neurons are also involved in parenting, infanticide, and allogrooming behavior (Chen et al., 2019; Hu et al., 2021; Raam and Hong, 2021; Wu et al., 2021).

A recent study (Baleisyte et al., 2022) confirmed our previous finding (Hong et al., 2014) that stimulation of MeApd GABAergic neurons using ChR2 induced a robust increase in aggression in male mice. Interestingly, however, they found that stimulation of these neurons using the original ChETA reduced rather than enhanced aggression, but this result was only obtained when high laser power and long duration photostimulation conditions were used. The authors suggested that this difference in results could potentially arise from differences in kinetics between the opsins, including a high plateau potential generated by ChR2 activation which may lead to depolarization block. If this conclusion were correct, it would imply a negative-rather than a positive-acting role for MeApd GABAergic neurons in aggression. Such a model would imply that inhibition of MeApd GABAergic neurons should promote, rather than inhibit, aggression. However, no loss-of-function experiments were performed by Baleiyste et al. (2022). To the contrary, as mentioned above inhibition or ablation of these neurons using multiple experimental modalities resulted in decreased, rather than increased aggression (Hong et al., 2014; Unger et al., 2015).

Taken together, these data make it unlikely that ChR2-induced aggression arises from depolarization block, i.e. inhibition of MeApd GABAergic neurons. Moreover, as depolarization block usually occurs on a longer time scale (15-30 s, Baleisyte et al., 2022), the rapid onset of induced aggression observed with ChR2 (∼1 s after the onset of stimulations, Hong et al., 2014) further argues against this possibility.

An alternative explanation for the difference between results obtained using ChR2 vs the original ChETA is that the latter has a substantially lower photocurrent than the former (Fig. 1B; Berndt et al., 2011; Mattis et al., 2012). Indeed, Baleisyte et al. (2022) also reported that ChETA photostimulation yielded substantially lower photocurrents than ChR2 in MeA GABAergic neurons *ex vivo*. This raises the possibility that the behavioral difference between the effects of the two opsins is due to the lower photocurrents evoked by ChETA, rather than to its faster kinetics, in comparison to ChR2. Weaker photocurrents would yield insufficient excitation to promote aggression and might promote competing behavioral drives (Hong et al., 2014; Lee et al., 2014).

To distinguish between these alternatives, we repeated the experiment using the improved ChETA_TC_, which has similar photocurrents as ChR2 but fast kinetics comparable to that of ChETA (Berndt et al., 2011; Gunaydin et al., 2010; Mattis et al., 2012). ChETA_TC_ does not produce high plateau depolarization and is therefore unlikely to cause depolarization block (Berndt et al., 2011; Mattis et al., 2012). We injected the MeApd of Vgat-Cre animals with AAV2-DIO-ChETA_TC_ (ChR2[ET/TC])-EYFP (Fig. 1C). We tested the injected animals using the resident-intruder assay by delivering 473 nm laser stimulation (15 s of 20 Hz, 20 ms stimulations, ∼5 mW/mm^2^). We found that optogenetic activation of MeApd Vgat^+^ neurons using ChETA_TC_ evoked robust aggressive behavior (Fig. 1D-E), while control Vgat-Cre animals injected with AAV2-DIO-EYFP and photostimulated showed no aggressive response (Fig. 1F). Together, these results suggest that an opsin with fast kinetics comparable to, but with larger photocurrents than, ChETA is capable of evoking aggression when actuated in MeApd Vgat^+^ neurons, similar to ChR2. Furthermore, because ChETA_TC_ does not produce high plateau depolarization and is therefore unlikely to cause depolarization block (Berndt et al., 2011; Mattis et al., 2012), this finding reduces the likelihood that the increased aggression produced by ChR2 arises from depolarization block of these neurons.

If depolarization block indeed explained the effect of optogenetic stimulation using ChR2 to promote aggression, the initiation of evoked attack should have a long latency relative to photostimulation onset because such a block requires longer timescales than optogenetically induced firing (15-30 s, Baleisyte et al., 2022). To test this, we examined the effect of short (2 sec) optogenetic stimulation trials using ChETA_TC_ (Fig. 2). To determine the shortest latency effect that could be obtained, stimulation trials were delivered when the animals were relatively close to each other (eliminating latency due to slow approach behavior). We found that a rapid onset of aggression could be robustly evoked in a majority of stimulation trials, whereas no increased aggression was associated with randomly intermingled sham (no laser) stimulations performed under identical conditions, i.e., when the animals were close to each other (Fig. 2). Strikingly, aggression could be evoked within 200 milliseconds in 26.2% of stimulation trials, and within 300 ms in 45.2% of stimulation trials. Such a short-latency behavioral effect further strengthens the conclusion that the increased aggression induced by ChR2 or ChETA_TC_ is unlikely to be due to depolarization block.

**Figure 2.**
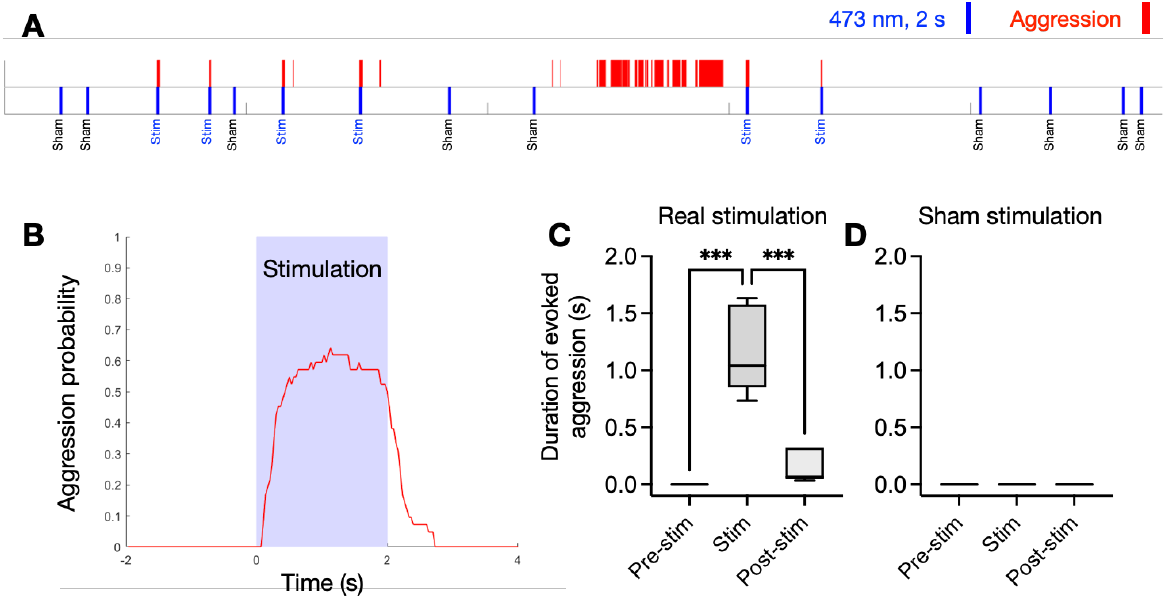
Short-duration stimulation using ChETA_TC_ induces rapid, robust aggression. **A**, Example behavioral raster plot including both sham and real stimulations. Aggression is induced by real stimulations but not sham. **B**, Probability of aggression during short-duration (2s) photoactivation of MeA GABAergic neurons expressing ChETA_TC_. Each real stimulation: 2 s of 20 Hz, 20 ms, 473 nm, ∼5 mW/mm^2^. **C, D**, Increased duration of aggression during photoactivation epochs (0–2s) relative to pre- (−2–0s) and post-stimulation (2–4s) epochs. **C**, Real stimulations; **D**, Sham stimulations. One-way ANOVA with post-hoc Tukey’s multiple comparisons. *** *p* < 0.001. n = 5 animals.

Given that ChETA_TC_ has fast kinetics similar to the original ChETA, yet produces similar behavioral phenotypes as ChR2, the different results obtained using the original ChETA are unlikely due to its faster kinetics as suggested by Baleisyte et al (2022). Rather, the most likely explanation for the difference is that ChETA has a weaker photocurrent than either ChR2 or ChETA_TC_. The reduction of aggression observed using ChETA may therefore be a consequence of a lower intensity of stimulation that can be achieved using this opsin, compared to ChETA_TC_ or ChR2. The reason for the inhibitory effect on aggression of low-intensity stimulation is not clear, and may involve promotion of a competing internal drive state (Hong et al., 2014) and/or the activation of a low-threshold subpopulation of inhibitory interneurons in MeApd. The latter is consistent with the observation that the MeA contains molecularly distinct GABAergic subpopulations (Chen et al., 2019; Wu et al., 2017). Achieving specific genetic access to these different subpopulations will allow further investigation of their role in aggression and other social behaviors.

It is noteworthy that the suppression of aggression observed by Baleisyte et al. (2022) using ChETA only occurred during repeated stimulation trials of long duration (30-60 s per trial for 5-12 trials) at high laser power (10-20 mW at the tip, which is 15-30 mW/mm^2^ irradiance), but not at lower laser power (Figure 1 and S1 in Baleisyte et al., 2022). These photostimulation conditions are considerably more intense than those used here with ChETA_TC_ (2 s at 5 mW/mm^2^), or previously with ChR2 (Hong et al., 2014; Lee et al., 2014; Lin et al., 2011). High laser power, long duration optogenetic stimulation conditions have been shown to cause thermal, electrophysiological, and behavioral artifacts, such as suppression of neural activity in multiple brain areas (Owen et al., 2019). Thus, the suppression of aggression observed by Baleisyte et al. (2022) may be a consequence of a suppression of MeA Vgat^+^ neuron activity caused by these artifacts, and should be interpreted with caution. Indeed, the fact that the results obtained using ChR2 or ChETA_TC_ are complementary to results from loss-of-function experiments using optogenetic inhibition, chemogenetic inhibition, and neuronal ablation (Hong et al., 2014; Unger et al., 2015) renders it more likely that those opsins, and not ChETA, mimic the natural role of MeApd Vgat^+^ neurons, i.e., to promote aggression.

Our results argue that concerns regarding possible depolarization block artifacts when using ChR2 in optogenetic stimulation experiments are better addressed using ChETA_TC_ rather than the original ChETA (Berndt et al., 2011; Mattis et al., 2012) to achieve faster opsin kinetics without sacrificing photocurrent magnitude. Moreover, given the consistent phenotype elicited by both ChR2 and ChETA_TC_ in our experiments, interpretations of circuit function from studies using ChR2 should not be dismissed or rejected by default, simply because of the opsin being used.

In conclusion, the present work, taken together with the results of Baleisyte et al. 2022, highlight the importance of taking into consideration the properties of the specific opsin tools used to probe the function of neural circuits. The continued development of improved opsins such as ChETA_TC_ affords closer approximations of the natural neuronal firing patterns in a cell type of interest, and will be crucial in aiding our understanding of circuit function.

## Acknowledgements

We thank Olexiy Kochubey and Ralf Schneggenburger (EPFL) for providing feedback on data described in this manuscript.

## Methods

### Experimental subjects and materials

Vgat-Cre knockin animals (Vong et al., 2011) were initially purchased from Jackson Laboratories and then backcrossed to the C57BL/6J background. Animals were housed and maintained on a reversed 12-h light-dark cycle for at least one week prior to stereotaxic surgery or behavioral testing. All the control experiments were done using animals with the same genetic background. Care and experimental manipulations of animals were in accordance with the National Institute of Health Guide for Care and Use of Laboratory Animals and approved by the Institutional Animal Care and Use Committee. AAV2-EF1a-DIO-ChR2[E123T/T159C]-EYFP (ChETA_TC_-EYFP) and AAV2-EF1a-DIO-EYFP were purchased from the University of North Carolina vector core facility.

### Behavioral assay for aggression

Aggression was examined using the resident-intruder assay (Blanchard et al., 2003). In this assay, an unfamiliar male (‘‘intruder’’) mouse was then introduced into the home cage of the tested resident, and then the resident and intruder were allowed to freely interact with each other. As in previous studies (Hong et al., 2014), the resident-intruder assay used in this study was designed to study offensive aggression. To avoid intruder-initiated aggression, a group-housed, nonaggressive C57BL/6J was used as the intruder male.

### Optogenetic stimulation

Vgat-Cre/+ mice at 8-10 weeks old were injected bilaterally into MeApd with an AAV expressing ChETA_TC_ or EYFP and bilaterally implanted with optic fibers (ferrule fiber). After a 3–4-week recovery period, the virus-injected animals were tested in the resident intruder assay. Before behavioral testing, a ferrule patch cord was coupled to the ferrule fiber implanted in the mouse using a zirconia split sleeve (Doric Lenses). Optic fibers were connected using an FC/PC adaptor (Doric Lenses) to a 473-nm blue laser (CNI Laser). Laser pulses were controlled through an Arduino micro-controller board and a customized MATLAB program. Blue (473 nm) light was delivered in 20 ms pulses at 20 Hz, at final output powers ∼5 mW/mm^2^.

## Notes

### Competing Interest Statement

The authors have declared no competing interest.

### Summary of Updates

This version of the manuscript has been revised to update the name of the improved ChETA.

